# Can patients with cerebellar disease switch learning mechanisms to reduce their adaptation deficits?

**DOI:** 10.1101/386466

**Authors:** Aaron L. Wong, Cherie L. Marvel, Jordan A. Taylor, John W. Krakauer

## Abstract

Systematic perturbations in motor adaptation tasks are primarily countered by learning from sensory-prediction errors, with secondary contributions from other learning processes. Despite the availability of these additional processes, particularly the use of explicit re-aiming to counteract observed target errors, patients with cerebellar degeneration are surprisingly unable to compensate for their sensory-prediction-error deficits by spontaneously switching to another learning mechanism. We hypothesized that if the nature of the task was changed – by allowing vision of the hand, which eliminates sensory-prediction errors – patients could be induced to preferentially adopt aiming strategies to solve visuomotor rotations. To test this, we first developed a novel visuomotor rotation paradigm that provides participants with vision of their hand in addition to the cursor, effectively setting the sensory-prediction-error signal to zero. We demonstrated in younger healthy controls that this promotes a switch to strategic re-aiming based on target errors. We then showed that with vision of the hand, patients with spinocerebellar ataxia could also switch to an aiming strategy in response to visuomotor rotations, performing similarly to age-matched participants (older controls). Moreover, patients could retrieve their learned aiming solution after vision of the hand was removed, and retain it for at least one year. Both patients and older controls, however, exhibited impaired overall adaptation performance compared to younger healthy controls (age, 18-33), likely due to age-related reductions in spatial and working memory. Moreover, patients failed to generalize, i.e., they were unable to adopt analogous aiming strategies in response to novel rotations, nor could they further improve their performance without vision of the hand. Hence, there appears to be an inescapable obligatory dependence on sensory-prediction-error-based learning – even when this system is impaired in patients with cerebellar degeneration. The persistence of sensory-prediction-error-based learning effectively suppresses a switch to target-error-based learning, which perhaps explains the unexpectedly poor performance by patients with spinocerebellar ataxia in visuomotor adaptation tasks.

## INTRODUCTION

Motor adaptation tasks investigate how systematic movement errors are reduced over repeated trials. In the laboratory, the experimenter introduces a perturbation that causes an unexpected movement error. Learning is largely thought to rely on sensory-prediction-errors (SPE), in which a mismatch between the intended and sensed location of the effector (e.g., a rotation or displacement of the cursor relative to the unseen hand) drives recalibration of the motor system. SPE-learning is an implicit process, is likely cerebellum-dependent (Bastian, 2006; Mazzoni and Krakauer, 2006; Wong and Shelhamer, 2011; Leow et al., 2017; Morehead et al., 2017), and appears to be the default means by which the motor system counters systematic errors (Shmuelof *et al.*, 2012; Vaswani *et al.*, 2015). However, recent studies have demonstrated that adaptation tasks can also be solved by reducing the error between the target and the cursor (target error, TE), such as by adopting explicit aiming strategies (Werner and Bock, 2007; Heuer and Hegele, 2008; Hegele and Heuer, 2010; Taylor and Ivry, 2011; Taylor *et al.*, 2014). In particular, TE-based learning contributes in visuomotor rotation tasks, in which individuals counter an imposed rotation of the cursor relative to the hand by aiming in a different direction than that of the target. In healthy individuals, SPE and TE-based learning sum together to allow rapid and sustained responses to the perturbation (Taylor *et al.*, 2014; Huberdeau *et al.*, 2015b). Disruption of one process leads to compensation by the other (e.g., greater TE-based strategy use compensates for an impaired SPE process) (Taylor et al., 2014; Bond and Taylor, 2015; Brudner et al., 2016; Schween and Hegele, 2017).

Patients with an impaired cerebellum have SPE learning deficits, leading to poor performance during motor adaptation tasks (Martin et al., 1996; Maschke et al., 2004; Smith and Shadmehr, 2005; Tseng et al., 2007). Why don’t these patients adopt an aiming strategy to compensate for their SPE-based learning deficit? Recent work has suggested one answer: TE-based learning may also be impaired in patients with spinocerebellar ataxia (Butcher et al., 2017), perhaps due to cognitive deficits (Schmahmann and Sherman, 1998; Suenaga *et al.*, 2008; Cooper *et al.*, 2010) that could hinder the development of re-aiming strategies during adaptation tasks (Anguera et al., 2010; Anguera et al., 2012; Christou et al., 2016). That said, when provided with an explicit aiming strategy, patients successfully overcome the visuomotor perturbation (Taylor et al., 2010). Hence patients may either be impaired at discovering aiming strategies, or TE-based learning mechanisms might be suppressed by SPE-based ones as we have previously suggested (Shmuelof *et al.*, 2012; Vaswani *et al.*, 2015), even though SPE-based learning is impaired in these patients.

To distinguish between these alternatives, we investigated whether we could teach individuals to switch to TE-based learning and rely less on their SPE-recalibration system. To do so, we provided individuals with real-time vision of their hand in addition to the cursor. This results in an SPE error-signal equal to zero because the hand is observed to move exactly as expected. Without a meaningful SPE error signal for learning, along with the additional information provided by seeing the hand, this paradigm should promote a spontaneous switch to TE-based aiming as long as patients with spinocerebellar ataxia retain the capacity to discover such strategies. As a supplemental analysis, we explored whether the use of TE-based strategies was correlated to an individual’s general cognitive abilities, specifically spatial and working memory capacity (Anguera et al., 2010; Anguera et al., 2012; Christou et al., 2016). Finally, we tested whether individuals could retain the use of such strategies when vision of the hand was subsequently removed, or if they could generalize when the rotation was in the opposite direction.

## METHODS

Three experiments were conducted. Experiment 1 compared how availability of vision of the hand changed the way that healthy individuals responded to a visuomotor rotation. Experiments 2 and 3 explored whether patients with spinocerebellar ataxia could use vision of the hand to improve their adaptation performance by promoting a switch to TE-based learning.

### Experimental Design

#### Participants

All participants were right handed and naïve to the purposes of the study. All individuals provided written informed consent to participate. Experimental methods were approved by the Johns Hopkins University School of Medicine institutional review board. As patients with spinocerebellar ataxia exhibit obvious kinematic impairments, experimenters were not blinded to participant diagnoses. Sample sizes were chosen to be consistent with previous studies (Tseng et al., 2007; Taylor et al., 2010; Butcher et al., 2017).

Thirty-four younger neurotypical participants (ages 18-34; average age, 23 years; 11 male) were recruited for Experiment 1. Participants were randomly assigned to one of two groups that differed in their exposure to vision of the hand during a visuomotor rotation paradigm. Five participants were removed after failing to develop a strategy to overcome the visuomotor rotation. This left 15 (alternating-vision) and 14 (all-vision) participants per group.

For Experiment 2, we recruited 15 patients with spinocerebellar ataxia (ages 47-82; average, 64 years; 11 female), 16 neurotypical age-matched older controls (ages 45-78; average 61 years; 9 female), and 16 neurotypical younger controls (ages 18-33; average 22 years; 9 female). All patients had a known genetic diagnosis or exhibited symptoms consistent with isolated cerebellar degeneration (no extrapyramidal signs, neuropathy, bradykinesia or parkinsonism, or history of strokes or cardiac problems). Patients had no cognitive impairments as noted by their neurologist. Two patients were excluded because they reported intentionally ignoring either their hand or the cursor when both were visible; the remaining patients are detailed in Table 1. These patients were well matched in age to the older controls (*t*-test, t(27) = 0.35, *p* = 0.72). One younger control participant was similarly excluded for intentionally ignoring the hand when it was visible.

**Table 1.** Patients with spinocerebellar ataxia

| Patient | Exp 2 | Exp 3 | Age | Sex | Diagnosis | ICARS | MOCA |
| --- | --- | --- | --- | --- | --- | --- | --- |
| P1 | Y | Y | 73 | F | SCA6 | 25 | 28 |
| P2 | Y | Y | 71 | F | SCA6 | 27 | 29 |
| P3 | Y | Y | 63 | F | SCA6 | 60 | 25 |
| P4 | Y | Y | 68 | M | ADCA | 8.5 | 27 |
| P5 | Y | Y | 63 | M | ADCA | 21 | 28 |
| P6 | Y | Y | 48 | F | SCA6 | 2 | 30 |
| P7 | Y | Y | 47 | F | SCA8 | 38 | 29 |
| P8 | Y | Y | 63 | F | ADCA | 29 | 29 |
| P9 | Y | Y | 53 | F | ADCA | 37 | 26 |
| P10 | Y | Y | 82 | M | ADCA | 23 | 29 |
| P11 | Y |  | 48 | M | SCA8 | 27 | 25 |
| P12 | Y |  | 67 | F | LOCA | 25 | 22 |
| P13 | Y |  | 72 | F | LOCA | 34 | 23 |
| P14 |  | Y | 56 | F | ADCA | 20 | 27 |
| P15 |  | Y | 56 | F | LOCA | 52 | 20 |

Patients that successfully completed Experiments 2 and/or 3 are shown in the table above.
Patients for which a clear genetic diagnosis of spinocerebellar ataxia (SCA) was not known could be subdivided into two groups. Patients with a known family history of cerebellar ataxia exhibiting an autosomal-dominant inheritance pattern are noted as ADCA; otherwise the diagnosis is reported as sporadic late-onset cerebellar ataxia (LOCA). In all cases, patients were screened for any cognitive deficits as noted by their treating neurologist.

Ten patients from Experiment 2 returned 10.8 ± 1.5 months later (range, 4-18 months) for Experiment 3. Two new patients were also recruited. Participants are listed in Table 1 (ages 47-82; average 61.76 years; 4 male). All patients reported that the two Experiments were similar, but only two explicitly recalled a strategy. These patients returned eleven and twelve months after completing Experiment 2; hence, time between sessions was unlikely to influence successful strategy retention.

### Experimental Paradigms

Participants made planar reaching movements by sliding their dominant (right) hand along a glass table. A cotton glove reduced friction between the hand and the table. Movement of the index finger was tracked using a Flock of Birds magnetic tracking system (Ascension Technology, VT, USA) at 130 Hz. An LCD monitor (60 Hz), reflected in a one-way mirror located directly above the table, displayed targets and a cursor in the plane of the hand (Fig. 1). A board could be placed beneath the semi-transparent mirror to obscure vision of the arm, leaving only the cursor and targets visible.

**Figure 1.**
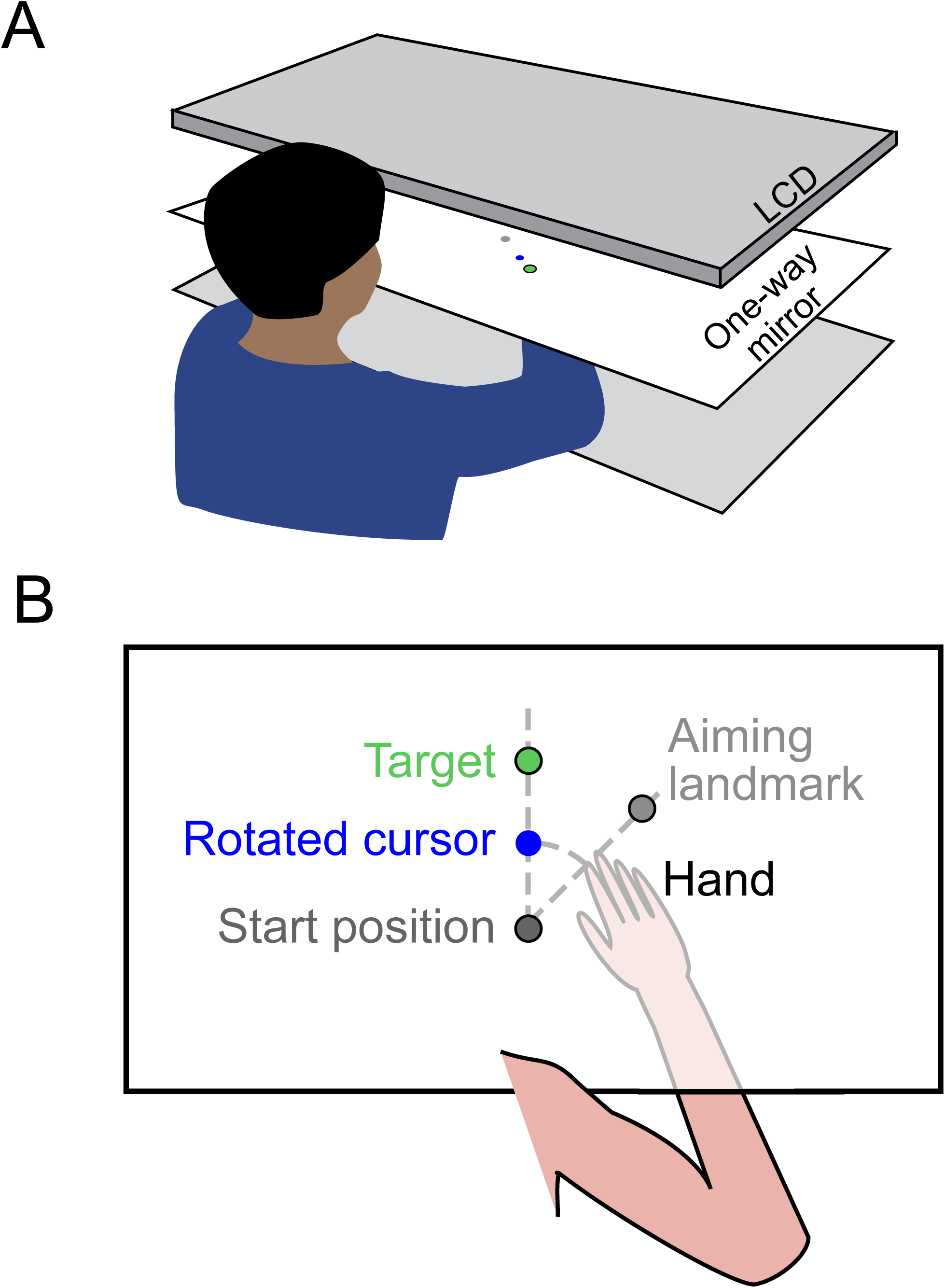
Experimental setup. (A) The experimental apparatus consisted of three tiers; an LCD monitor, a one-way mirror in the middle, and a table surface at the bottom. (B) Top-down view of what participants saw during an experiment. Stimuli (targets and the cursor) were always visible by observing the reflection of the LCD monitor in the mirror. In contrast, by adjusting the relative light levels below and above the one-way mirror, the hand, located on the surface of the table under the mirror, could be made visible or hidden from the participant’s view when looking through the mirror. When the visuomotor rotation was imposed, the cursor and the hand could be observed to diverge as the hand moved away from the start position.

Targets could appear at any of four positions equally spaced along a circle of radius 8 cm from the starting position. Visible landmarks were spaced 45° apart and could be used as aiming targets, although they were never described as such to the participants. On each trial, participants made a rapid shooting movement through the target. The cursor was only visible during the outward reach; to facilitate returning to the starting position it was replaced by a ring whose diameter reflected the magnitude of the hand from the center. For Experiments 2 and 3, a somatosensory landmark (raised bump on the table) also indicated the start position. After each experiment, participants completed a questionnaire to assess how they approached the task.

#### Experiment 1

In Experiment 1, younger healthy participants completed five blocks of trials (Fig. 2A). In block 1 (baseline, 60 trials), participants saw only the non-rotated cursor. In the next block (rotation block 1), all participants were given vision of their hand along with the cursor, and completed 20 baseline trials followed by 80 trials in which the cursor was rotated 45° counterclockwise relative to the hand. To overcome the rotation, participants had to aim 45° clockwise (toward the next landmark); although no explicit instructions were given, participants were expected to discover this strategy since they could see their hand and the cursor. During the second rotation block (60 trials), vision of the hand was removed for one of the two groups (alternating-vision group). For both groups, participants were instructed to continue doing whatever they had done previously; i.e., continue aiming the same as in the prior block. In the third rotation block (60 trials), both groups could again see their hand. Finally, there was a washout block of 80 non-rotated trials without vision of the hand.

**Figure 2.**
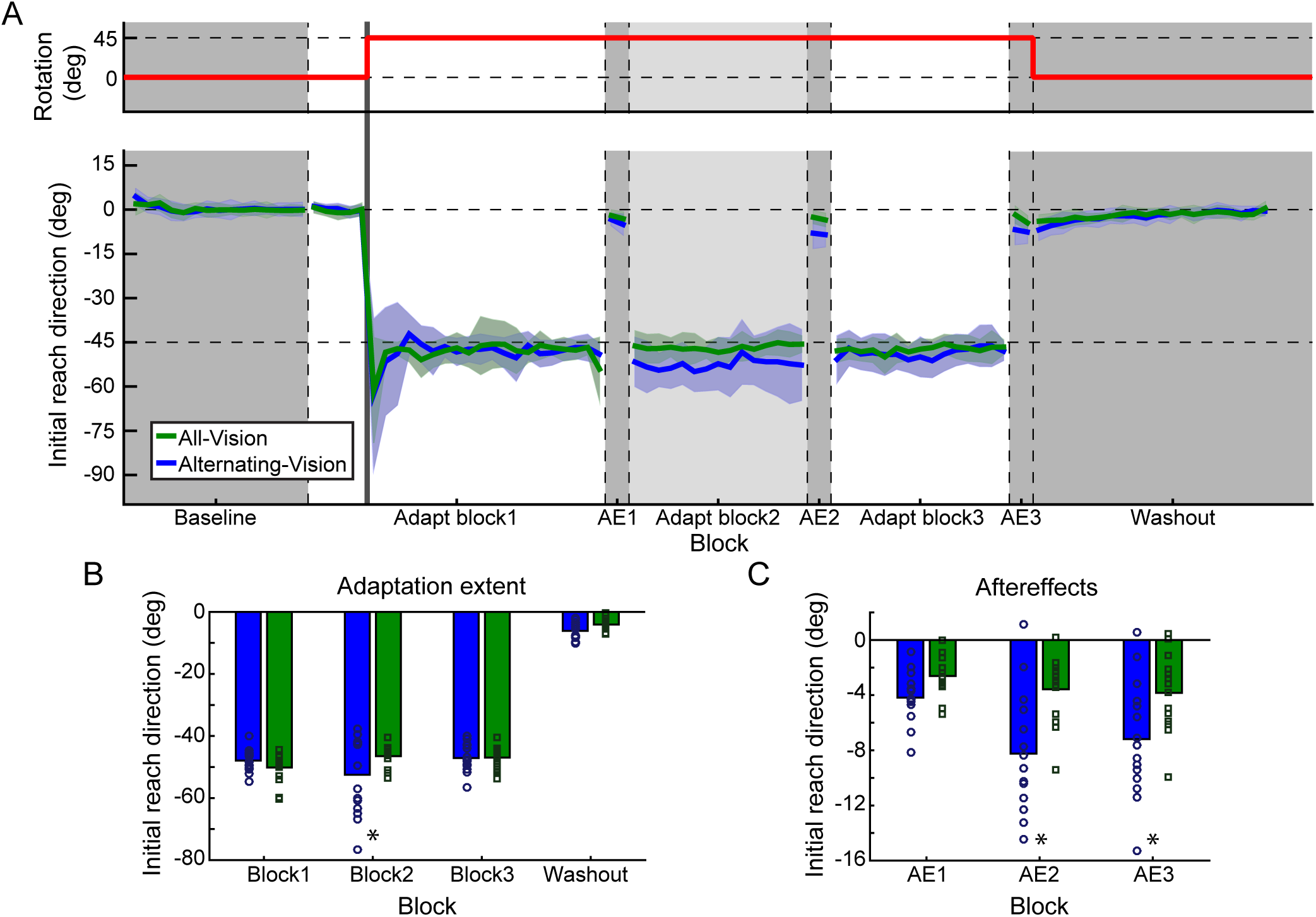
Experiment 1. (A) Top panel: the thick red line represents the direction in which the cursor has been rotated relative to the position of the hand. Bottom panel: Initial reach direction across the three adaptation blocks for the two groups. Dark gray sections are times when the hand is not visible to both groups; during the light gray section, the hand is hidden from the Alternating-Vision group (blue) but remains visible to the All-Vision group (green). During the three aftereffect (AE) blocks, neither vision of the hand nor the cursor is available to participants. (B) Summary of the average initial reach direction measured during the last two cycles of each adaptation block and the first two cycles of the washout block, for the All-Vision (green) and Alternating-Vision (blue) groups. Significant differences between groups are denoted by an asterisk. (C) Average reach direction during each of the three aftereffect blocks (average of both cycles) for the two groups as in (B).

After each rotation block, participants were given a brief aftereffect (AE) assay to assess the current adaptation state in the absence of any strategy use. Participants made 8 reaches without vision of the hand or the cursor, and were asked to stop using any strategies and instead aim directly for the target.

#### Experiments 2 and 3

Experiment 2 (Fig. 3) consisted of a baseline block (60 trials), followed by three consecutive learning bouts. Each bout involved two blocks: (1) eight baseline trials followed by 48 rotation trials, and (2) eight rotation trials followed by 48 washout trials. The visuomotor rotation direction was clockwise in Bout 1 and counter-clockwise in Bouts 2 and 3. Vision of the hand was available only during Bout 2. This experiment examined whether individuals could adopt TE-based learning strategies with vision of the hand (Bout 2), and re-apply those strategies when encountering the same rotation without vision of the hand (Bout 3).

**Figure 3.**
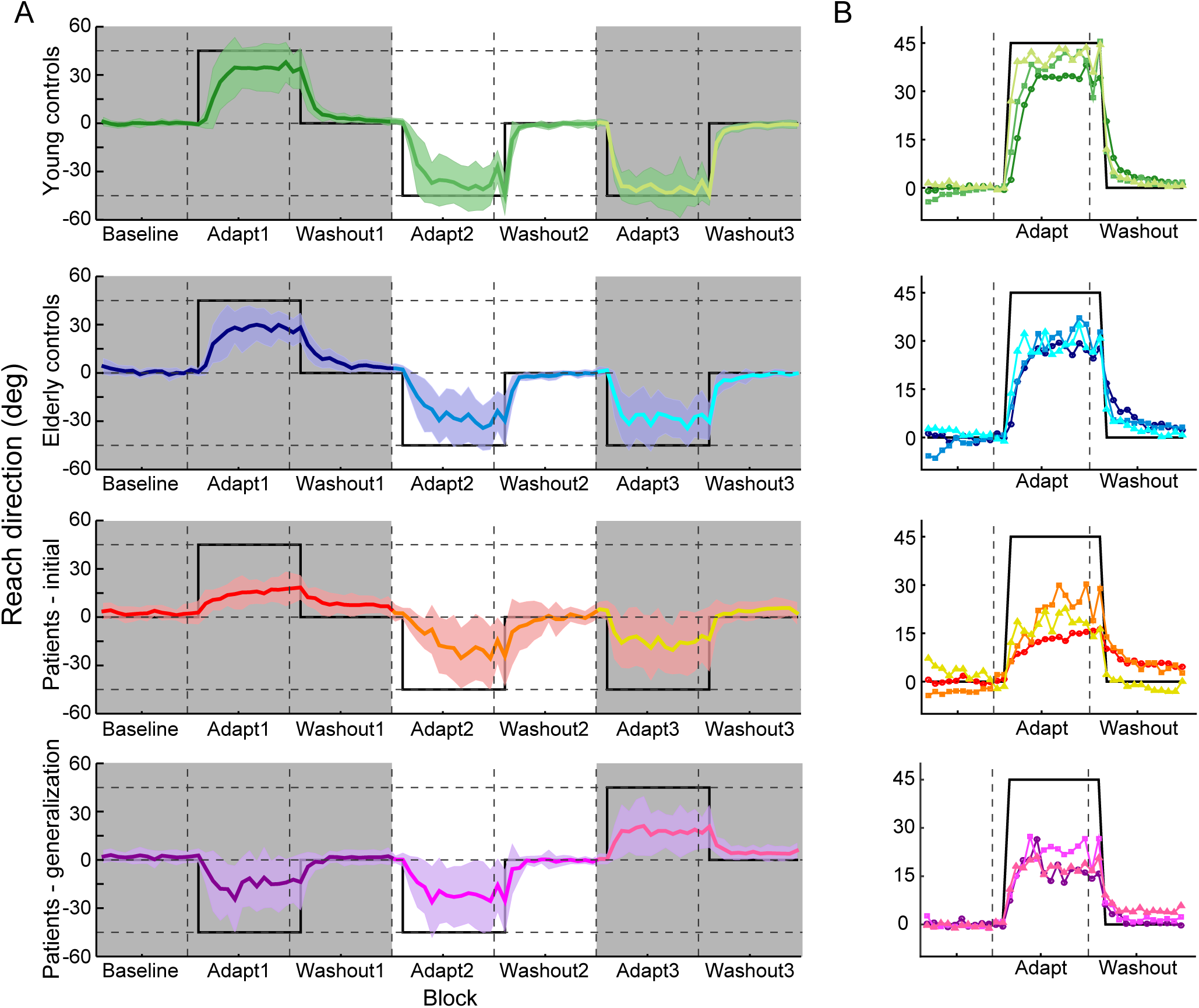
Experiments 2 and 3. (A) Initial reach direction throughout the three adaptation bouts for younger controls in Experiment 2 (top panel), old controls in Experiment 2 (second panel from the top), patients with spinocerebellar ataxia in Experiment 2 (third panel), and patients in Experiment 3 that examined generalization (bottom panel). Dark shaded regions represent times when the hand was not visible to the participant (i.e., Bouts 1 and 3). The thick black line represents the reach direction required to completely counteract the visuomotor rotation. (B) Overlays of initial reach direction from each adaptation bout; panels correspond to those in Figure 3A. Data have been normalized within each subject to the 4 cycles immediately preceding the onset of the visuomotor rotation to compare the initial adaptation rate and the total adaptation extent. Bouts are color-coded, with darker colors occurring earlier in the session and lighter colors occurring later. Thick black lines reflect the ideal reach direction to counteract the visuomotor rotation.

Motor impairments were assessed using the International Cooperative Ataxia Rating Scale (ICARS) (Trouillas et al., 1997). Patients and older controls also completed a cognitive test battery (see Supplementary Material).

Experiment 3 (Fig. 3) was identical to Experiment 2 except for the order of experienced rotations; the rotation was counter-clockwise in Bouts 1 and 2, and clockwise in Bout 3. As before, vision of the hand was only available during Bout 2. This experiment enabled us to assess long-term retention (comparing Experiment 2, Bout 3 to Experiment 3, Bout 1) as well as generalization of TE-strategies to a rotation in the opposite direction (comparing Bouts 2 and 3 in Experiment 3).

### Data Analysis

Data were analyzed offline using MATLAB (The MathWorks, Natick, MA). Reaches were selected according to a velocity criterion (tangential velocity >0.05 m/s in Experiment 1, or >0.03 m/s in Experiments 2 and 3) and verified by visual inspection. Reach velocity was computed as the derivative of hand position after smoothing with a second order Savitzky-Golay filter (frame size, 19 samples). Reach direction was defined as the direction of the velocity vector, relative to a straight line between the starting and target locations, 100 ms (Experiment 1) or 250 ms (Experiment 2, since reaches were typically slower) after movement onset (positive angles are counter-clockwise). Reach angles greater than a threshold (Experiment 1, 45°; Experiments 2 and 3, 60°) were excluded as aiming errors.

Every four reaches were averaged into cycles. Learning was quantified as the average reach angle of the last two (Experiment 1) or four (Experiments 2 and 3) cycles prior to the block break (to exclude forgetting due to the break; Smith *et al.*, 2006; Joiner and Smith, 2008). In Experiment 1, the average of all 8 trials for each aftereffect assay was also examined. In Experiments 2 and 3, learning curves were normalized by subtracting the average reach direction of the two baseline cycles just prior to rotation onset to account for any incomplete washout. Savings was quantified as the initial adaptation rate (average of the first four cycles following rotation onset).

Reaction time was calculated as the time between target onset and movement onset. Movement time was defined as the time from movement onset to when the hand exceeded 8 cm from the starting position.

#### Statistical Analyses

Reach directions were compared with mixed-effects models in R using the *lme4* package (Bates et al., 2015) with fixed effects of Group and Bout and random effects of Participant. Significance was determined using likelihood ratio tests comparing models with and without the factor of interest; post-hoc tests were performed using the generalized linear hypothesis testing function in the *multcomp* package (Hothorn et al., 2008) and adjusted for multiple comparisons using Bonferroni-Holm corrections. In all cases, S.E.M. is reported. For Experiment 2, stepwise regressions were also performed to examine the relationship between learning extent and motor and cognitive assessments in the patient and older control groups (see Supplementary Material).

## RESULTS

### Vision of the hand switched learning from sensory prediction errors to target errors

In Experiment 1, we hypothesized that allowing individuals to observe both their actual hand and the cursor would induce a switch to a TE-based strategy to overcome a visuomotor rotation. This is because the SPE signal would be equal to zero (i.e., the observed hand moves exactly as expected according to the issued motor command). To observe the impact of vision of the hand, we compared participants who saw their hand throughout the entire experiment to those for which vision of the hand was obscured during the second of three rotation blocks. We will first examine performance during learning and washout, then discuss the aftereffect trials.

Providing participants with vision of the hand (Fig. 2A,B) had no effect on baseline performance (reach direction, χ^2^(1) = 1.96, *p* = 0.16; latency, χ^2^(1) = 0.12, *p* = 0.74; movement time, χ^2^(1) = 0.88, *p* = 0.35). However, there was a significant effect of group (χ^2^(4) = 15.80, *p* = 0.003) and block (χ^2^(4) = 571.17, *p* < 0.001), as well as a significant interaction (χ^2^(4) = 17.74, *p* = 0.001). The effect of block was primarily driven by the initial response to counteract the visuomotor rotation (Fig. 2; baseline, 0.09° ± 0.36°; learning extent in block 1, −48.97° ± 1.17°; difference between blocks, *z* = 50.22, *p* < 0.001; no difference between groups, *z* = 1.43, *p* = 0.69). Because the hand was visible when the visuomotor rotation was introduced, the typical exponential learning curve associated with adaptation experiments was not observed; within a few trials participants immediately understood the perturbation and countered it fully by reaiming. About 80% of participants reported aiming one target clockwise in response to the TE due to the discrepancy between the cursor and the hand.

The group effect was largely driven by a difference in reach direction during the second rotation block when vision of the hand was removed in one group(all-vision group, −46.50° ± 0.93°; alternating-vision group, −52.49° ± 3.18°, *z* = −3.84, *p* = 0.001). Removing vision of the hand resulted in drift away from the target, which has previously been associated with implicit SPE-based learning (Mazzoni and Krakauer, 2006). Note that here we do not see a gradual increase in SPE-induced drift, likely because between-participants performance was quite variable during this block as individuals had learned to continuously re-adjust their aiming direction in response to errors (with varying degrees of success) rather than adopt a fixed aiming strategy. Restoration of vision in the third rotation block enabled individuals in the alternating-vision group to re-adjust their aim using TE, again resulting in similar performance between groups (all-vision group, −46.94° ± 0.95°; alternating-vision group, −47.08° ± 1.13°; *z* = −0.09, *p* = 1.00). Finally, during the final washout block performance remained similar across groups (all-vision group, −4.08° ± 0.50°; alternating-vision group, −6.12° ± 0.77°, *z* = −1.31, *p* = 0.61). No pairwise differences were observed in latency or movement time for any block (p > 0.11).

Vision of the hand facilitated consistent success at hitting the target. This could have arisen from one of two reasons. Vision of the hand may have simply enabled participants to overcome any underlying drift by continuously applying small TE-based adjustments, giving the appearance of stable performance. Alternatively, vision of the hand may have interfered with SPE-based learning such that no drift occurred when vision was available. To distinguish between these hypotheses, we used aftereffect trials to evaluate the adaptation state of the motor system after each rotation block. In these trials, participants had no feedback of the hand or the cursor and were asked to aim directly for the target; thus, any deviation in reach direction directly reflected the participant’s adaptation state.

Measured aftereffects (Fig. 2C) also exhibited a significant effect of group (χ^2^(2) = 11.66, *p* = 0.003) and block (χ^2^(2) = 36.58, *p* < 0.001), as well as a significant interaction (χ^2^(2) = 10.60, *p* = 0.005). Aftereffects immediately following the initial rotation block were small and consistent between the two groups (all-vision group, −2.61° ± 0.39°; alternating-vision group, −4.17° ± 0.46°; *z* = −1.44, *p* = 0.45). However, aftereffects were much larger upon removal of vision in the second rotation block (all-vision group, −3.57° ± 0.65°; alternating-vision group, - 8.24° ± 1.11°; *z* = −4.30, *p* < 0.001), suggesting that the non-zero SPE signal induced drift when vision of the hand was unavailable. The magnitude of this aftereffect is consistent with previous work suggesting SPE-based drift of 10°-15° (Morehead *et al.*, 2017; Kim *et al.*, 2018).

Surprisingly, after the third rotation block when vision of the hand was restored, the aftereffects were similar to those following the second rotation block for both groups (all-vision group, −3.83° ± 0.76°; alternating-vision group, −7.19° ± 1.07 °; difference in aftereffects between blocks 2 and 3: all-vision group, *z* = 0.39, *p* = 0.81, alternating-vision group, *z* = −1.60, *p* = 0.44; difference in aftereffects for block 3 between groups, *z* = −3.09, *p* = 0.01). Thus, although participants in the alternating-vision group appeared to counteract the drift during the third rotation block (Fig. 2B), this was not a true washout of SPE-based learning: the persistent larger aftereffect observed for this group suggests that the SPE-based drift was instead overcome by applying TE-based aiming without changing the current adapted state (i.e., providing vision of the hand “froze” the adapted state). Taken together, these findings suggest that vision of the hand prevented any SPE-based changes during learning or washout. That is, the state of the motor system as assessed by aftereffect trials did not change when vision of the hand was available: setting SPE to zero by providing vision of the hand preventing any modification of the current adapted state, including any forgetting or washout.

### Vision of the hand modified performance in response to a visuomotor rotation

The findings from Experiment 1 suggest that vision of the hand serves two purposes. First, it sets SPE to zero because the hand is seen to move exactly as expected, preventing SPE-based learning from modifying the current adaptation state of the limb. Second, by providing feedback of both the hand and cursor relative to the target, it encourages the use of compensatory TE-based aiming strategies. Thus, we hypothesized that if patients with spinocerebellar ataxia could see their hand, this would unmask their ability to use TE-based strategies to overcome the rotation. This might then encourage the use of TE-based strategies in future adaptation tasks, particularly for patients with cerebellar damage for which SPE-based learning is impaired.

Therefore, in Experiment 2 we examined how patients with spinocerebellar ataxia responded to a visuomotor rotation with and without vision of the hand (Fig. 3). Patients completed three short bouts: the first bout assayed naïve learning of a visuomotor rotation without vision of the hand, the second provided vision of the hand to encourage adoption of a TE-based strategy, and the third tested retention of this strategy when encountering the same perturbation without vision of the hand. Performance was compared to a group of age-matched older controls and a group of younger controls (Table 2). To test for spontaneous adoption of aiming strategies without prompting due to a change in instructions, no aftereffect trials were introduced in this experiment; this also kept the experimental design as similar as possible to previous studies that have examined adaptation in patients with spinocerebellar ataxia.

**Table 2:** Reach Direction at Baseline and at the end of Adaptation

|  | Baseline | Bout 1 | Bout 2 | Bout 3 |
| --- | --- | --- | --- | --- |
| Younger Controls | $0.20^{\circ} \pm 0.21^{\circ}$ | $37.25^{\circ} \pm 3.17^{\circ}$ | $42.00^{\circ} \pm 3.06^{\circ}$ | $43.18^{\circ} \pm 2.42^{\circ}$ |
| Older Controls | $0.58^{\circ} \pm 0.39^{\circ}$ | $26.97^{\circ} \pm 1.99^{\circ}$ | $33.07^{\circ} \pm 2.88^{\circ}$ | $31.03^{\circ} \pm 2.44^{\circ}$ |
| Patients-Exp2 | $2.11^{\circ} \pm 1.32^{\circ}$ | $14.02^{\circ} \pm 2.50^{\circ}$ | $24.40^{\circ} \pm 5.07^{\circ}$ | $20.65^{\circ} \pm 4.45^{\circ}$ |
| Patients-Exp3 | $2.40^{\circ} \pm 0.87^{\circ}$ | $18.01^{\circ} \pm 3.94^{\circ}$ | $25.27^{\circ} \pm 5.32^{\circ}$ | $18.99^{\circ} \pm 4.08^{\circ}$ |

At baseline, younger controls were significantly more accurate than patients with regard to their initial reach direction (Table 2; post-hoc *t*-test, *p* = 0.03); all other pairwise comparisons between groups were not significant (patients versus older controls, *p* = 0.17; older controls versus younger controls, *p* = 0.63). Upon exposure to the visuomotor rotation, we observed a significant group difference in the extent to which individuals adapted throughout the session (χ^2^(2) = 77.35, *p* < 0.001) as well as a significant effect of bout (χ^2^(2) = 32.73, *p* < 0.001), although there was no significant interaction between group and bout (χ^2^(2) = 3.93, *p* = 0.41). To examine these differences in greater detail, we will first summarize differences between the patient and older control groups, then contrast this against the younger controls.

#### Vision of the hand improved performance for patients with spinocerebellar ataxia

During initial exposure to the visuomotor rotation without vision of the hand (Bout 1), patients exhibited impaired learning compared to older controls (Table 2; post-hoc test, *z* = 3.58, *p* = 0.004). This is not surprising and has been reported many times previously (Martin *et al.*, 1996; Maschke *et al.*, 2004; Smith and Shadmehr, 2005; Tseng *et al.*, 2007; Butcher *et al.*, 2017).

Providing vision of the hand in Bout 2 significantly improved the response to the visuomotor rotation compared to Bout 1 (Table 2) for patients (*z* = 4.32, *p* < 0.001) and older controls (*z* = 3.03, *p* = 0.02). This improvement likely reflected the adoption of TE-based strategies (consistent with this hypothesis, performance was largely limited by visuospatial memory capacity - Supplementary Results). In fact, patients exhibited greater performance improvements from Bouts 1 to 2 (by 10.38° ± 5.79°) compared to older controls (6.10° ± 2.85°). Thus although patients on average still compensated for the rotation a little worse than controls, a significant difference was no longer observed across groups in this Bout (*z* = 2.55, *p* = 0.07).

Finally, in Bout 3 vision of the hand was removed, but there was no further change in performance compared to Bout 2 (Table 2). Although patient performance decreased somewhat in Bout 3, this change was not significant (*z* = 1.17, *p* = 0.96) and remained better than that exhibited during initial exposure to the rotation (*z* = 3.10, *p* = 0.02). Thus whatever patients learned in the second bout was likely retained (but not further improved) even after vision of the hand was removed. Similarly, the performance exhibited by older controls also remained consistent during Bout 3 compared to Bout 2 (*z* = 1.08, p = 0.96). Although older controls again performed better than patients, this group difference did not reach significance (*z* = 2.68, *p* = 0.06). Thus in general, by requiring participants to stop using SPE-based learning and adopt TE-based aiming strategies, vision of the hand led to an improvement in response to the visuomotor rotation that was retained after vision of the hand was removed.

Patients and older controls also exhibited savings (larger performance gains in the first 4 cycles after rotation onset) in Bout 3 compared to Bout 2 (patients: *z* = −2.09, *p* = 0.07; controls: *z* = −2.46, *p* = 0.03) but not between Bouts 1 and 2 (patients: *z* = 1.12, *p* = 0.26; controls: *z* = 0.80, *p* = 0.42). This savings without an accompanying performance improvement in Bout 3 suggests that the learned aiming direction in Bout 2 was simply retrieved in Bout 3, consistent with previous work attributing savings to recall of an aiming strategy (Haith *et al.*, 2015; Huberdeau *et al.*, 2015a; Morehead *et al.*, 2015).

#### Younger controls consistently outperformed patients and older controls

Although patients with spinocerebellar ataxia improved their asymptotic performance to a level comparable to that of older controls, neither group achieved perfect compensation for the visuomotor rotation. One possibility is that participants didn’t have enough exposure to the rotation; this seems unlikely given that performance reached an asymptote and did not improve further in Bout 3. Alternatively, the change in rotation direction between Bouts 1 and 2 could have led to interference that blunted performance in the subsequent adaptation bouts (Krakauer *et al.*, 2005). To address this concern, we examined the behavior of a group of younger control participants in this same task.

Throughout the task, younger controls responded to the visuomotor rotation by changing their reach direction to almost completely compensate for the 45° rotation, exceeding that of both the patient and older control groups (Table 2). This difference in performance was already evident in the first adaptation bout (younger controls versus patients, *z* = 6.26, *p* < 0.001; younger controls versus older controls, *z* = 4.63, *p* < 0.001). Younger controls also outperformed the other groups in Bout 2 when vision of the hand became available (younger controls versus patients, *z* = 4.80, *p* < 0.001; younger controls versus older controls, *z* = 3.88, *p* = 0.001), and in Bout 3 when vision of the hand was again removed (younger controls versus patients, *z* = 5.85, *p* < 0.001; younger controls versus older controls, *z* = 5.51, *p* < 0.001).

Thus younger controls exhibited consistent performance across all three bouts. In particular, when vision of the hand was provided, younger controls exhibited only modest improvements in performance (average change in reach direction, 4.75° ± 2.89°, *z* = 2.22, *p* = 0.16), although this could be a ceiling effect. When vision was subsequently removed, the performance of younger controls also remained consistent; (average change in reach direction, 1.18° ± 3.02°, *z* = 0.55, *p* = 0.96). This suggests that both older controls and patients were unable to achieve the performance exhibited by younger controls not because of something unusual about the paradigm, but instead simply because they were older.

Younger controls became successively faster at responding to the perturbation each time it occurred (pairwise comparisons in learning rate between Bouts, *p* < 0.014). From Bout 1 to Bout 2 this could have been the result of savings, such as through use-dependent learning of successful actions (Huang *et al.*, 2011), or simply the result of adopting a completely explicit reaiming strategy as in Exp 1. Savings from Bout 2 to 3, on the other hand, likely arose from retrieval of the previously learned solution to the rotation (Haith *et al.*, 2015; Huberdeau *et al.*, 2015a; Morehead *et al.*, 2015).

### Target-error-based strategies were rotation-specific

Patients responded to the visuomotor rotation better when they had previously been able to see their hand, but it was unclear if they were recalling something about the specific rotation or had learned to apply a more general TE-based strategy for any rotation. To test this, in Experiment 3 the rotations when vision of the hand was available (Bout 2) and when vision of the hand was subsequently removed (Bout 3) were in opposite directions (Fig. 3).

We observed a significant main effect of bout (χ^2^(1) = 12.09, *p* = 0.002). Pairwise posthoc testing revealed a significant increase in the extent of learning achieved from Bout 1 to Bout 2 (Table 2; *z* = 3.29, *p* = 0.002) but a significant decrease in performance between Bouts 2 and 3 (Table 2; *z* = 2.83, *p* = 0.009) and no difference between Bouts 1 and 3 (*z* = 0.47, *p* = 0.64). Hence, patients could not generalize their aiming strategies to a novel rotation. Instead, this is consistent with the argument that in Bout 3 of Experiment 2, patients were simply retrieving the aiming direction they learned in Bout 2 rather than applying a general aiming strategy.

We also observed a significant retention in performance for the 10 patients that participated in both Experiments 2 and 3, despite having approximately a year between testing sessions. Performance in the first bout of Experiment 3 was better than in the first bout of Experiment 2 (χ^2^(1) = 16.11, *p* < 0.001), and comparable to the last bout in Experiment 2 (χ^2^(1) = 2.44, *p* = 0.12). Interestingly, only two patients were able to explicitly recall the aiming strategy learned in the previous Experiment, suggesting that whatever was being retrieved was not purely explicit but may have become automatized (Huberdeau *et al.*, 2017) or cached (Haith and Krakauer, 2018; McDougle and Taylor, 2018). These findings are consistent with the idea that patients could retrieve an aiming solution they were taught, but could not develop a general reaiming strategy.

## DISCUSSION

In these studies, we sought to understand why patients with spinocerebellar ataxia do not spontaneously adopt TE-based aiming strategies to compensate for their SPE-learning impairments. By providing direct vision of the hand, we effectively set the SPE signal to zero (at least in healthy individuals), changing how individuals responded to the visuomotor rotation perturbation. In Experiment 1, we found that in healthy younger adults, vision of the hand effectively encouraged a switch to TE-based strategies to overcome the rotation. According to post-test questionnaires these strategies were largely explicit and focused on re-aiming the hand to a different location. Equally important, by setting SPE to zero we effectively froze the current adapted state of the motor system and not only prevented any further learning, but also any washout or forgetting. This is consistent with prior work showing an implicit learning component that accounts for 10°-15° of overall learning (Morehead *et al.*, 2017; Kim *et al.*, 2018) and is very stable despite attempts to wash it out (Joiner and Smith, 2008).

In Experiment 2, we reaffirmed that patients with spinocerebellar ataxia were impaired at compensating for a visuomotor rotation. Notably, however, when given vision of the hand, they could develop and apply aiming strategies to significantly improve their performance. Thus we unmasked a latent capacity for re-aiming that was not evident during a traditional visuomotor rotation assay. Age-matched controls also benefited when they could see their hand, but remained unable to match the performance of healthy younger controls. Nevertheless, all individuals could retrieve that aiming solution when vision of the hand was subsequently removed. However, lack of further improvement beyond that exhibited when vision of the hand was available (Experiment 2), along with the lack of generalization to a rotation in the opposite direction (Experiment 3), suggest that individuals were not learning to approach the task differently (i.e., meta-learning about aiming). Instead, they simply recalled the aiming solution learned when vision of the hand was available. Together, these findings suggest that patients with spinocerebellar ataxia actually face two distinct challenges: a cerebellar-damage-associated deficit in SPE-based learning that prevents patients from adapting comparably to age-matched controls, and a more general, age-related deficit that prevents older adults from performing as well as younger controls.

Our results reveal that patients with spinocerebellar ataxia have a latent capacity to employ TE-based aiming strategies that is not invoked under typical circumstances. Instead, there is a strong bias toward SPE-based learning, despite an impairment in this process. In other words, various learning processes appear to be in competition, with SPE-based learning typically winning out over and partially suppressing other processes such as reward-or TE-based learning. This has also been suggested to occur in healthy individuals (Shmuelof *et al.*, 2012; Vaswani *et al.*, 2015; Cashaback *et al.*, 2017). Rather than spontaneously increasing the contribution of TE-based strategies to compensate for SPE-based learning deficits, however, patients with spinocerebellar ataxia instead exhibited poor overall learning. Moreover, when SPE-errors were restored in Bout 3, both patients and controls rapidly retrieved the TE-based aiming solution they had previously learned but did not appear to modify it to further improve performance. Thus a nonzero SPE prevents the use of a purely TE-based approach to overcome a perturbation. This seems true not for patients but also older controls, who in Bout 3 of Experiment 2 also retrieved but did not improve upon the aiming solution they acquired in Bout 2. Hence, SPE-based learning appears to be an obligatory default learning mechanism that is used preferentially instead of TE-based learning.

As for the use of TE-based strategies, there appear to be two distinct limits on how well patients can perform. The first is that developing and using aiming strategies seems limited by cognitive resources like visuospatial and working memory. Our supplementary data suggest that patients with greater visuospatial and working memory capacity were better at developing and retaining TE-based aiming solutions. The second potential source of impairment in the use of TE-based strategies can be seen in older individuals in general. Regardless of diagnosis, no older participant was able to find a TE-based strategy that fully compensated for the visuomotor rotation. Most either reported that there was no systematic perturbation, or adopted a target-specific strategy; only three patients and one control framed their response strategy as a clockwise or counterclockwise rotation. In contrast, about 80% of younger adults in Experiment 1 and 80% in Experiment 2 described their strategy in that manner. Indeed, a few older participants complained that they recognized how they should have aimed on one trial (e.g., “to the right”) but became confused on the next trial because the target was different and required aiming in a different direction (e.g., “up”). As age impacts working memory and processing speed (for review see Reuter-Lorenz and Sylvester, 2005) as well as the use of strategies in problem-solving (for review, see Lemaire, 2010), it is not surprising that older participants exhibited worse performance compared to younger adults (McNay and Willingham, 1998; Buch *et al.*, 2003; Bock and Girgenrath, 2006; Seidler, 2006; Heuer and Hegele, 2008; Hegele and Heuer, 2013; Trewartha *et al.*, 2014; Huang *et al.*, 2017; Vandevoorde and Orban de Xivry, 2018; Wolpe *et al.*, 2018). A link between spatial working memory and adaptation tasks has even been reported in younger healthy individuals (Anguera et al., 2010; Anguera et al., 2012; Christou et al., 2016), implying that in general performance during adaptation tasks may be influenced by visuospatial memory capacity.

Thus patients with spinocerebellar ataxia are faced with two distinct challenges that should not be conflated. First, they have a deficit in SPE-based learning for maintaining a calibrated motor system (Weiner et al., 1983; Martin et al., 1996; Tseng et al., 2007; Rabe et al., 2009; Criscimagna-Hemminger et al., 2010). Second, these patients are older, and age has been previously associated with a general impairment in the ability to use cognitive strategies to compensate for perturbations (Uresti-Cabrera *et al.*, 2015; Vandevoorde and Orban de Xivry, 2018; Wolpe *et al.*, 2018). These cognitive abilities might be further stressed by cerebellar impairment: patients with spinocerebellar ataxia have been reported to have cognitive impairments relative to age-matched controls (Schmahmann and Sherman, 1998; Mandolesi et al., 2001; Cooper et al., 2010; Hogan et al., 2011; Butcher et al., 2017), although it is unclear if this arises directly from cerebellar degeneration or is an indirect result of needing additional cognitive resources to support motor processes that in healthy individuals are fairly automatic. Regardless, this decreases the likelihood of adopting aiming strategies *de novo*, despite an intact capacity to do so as we demonstrated in Experiment 2. On top of this cognitive impairment, it appears that the availability of SPE-based learning may suppress the use of other learning systems, at least to some extent. (The preferential use of one learning mechanism and suppression of an alternative has also been observed in other systems (Brembs, 2009; Anderson *et al.*, 2011)). The sum of these deficits accounts for the performance impairments seen in patients with cerebellar degeneration during adaptation tasks. One important clinical implication of these findings is that patients with spinocerebellar ataxia may actually be able to compensate for their motor impairments using strategic approaches if the inherent reliance on SPE-based learning can be circumvented.

In summary, we have demonstrated that individuals cannot avoid SPE-based learning mechanisms during adaptation tasks, even when those mechanisms are impaired. Only when the SPE signal is set to zero – such as by providing vision of the hand – can individuals solve this task by turning completely to alternative TE-based approaches that rely on cognitive processes. Performance improvements learned in this manner can be subsequently retrieved to boost performance both in healthy older adults as well as in patients with cerebellar degeneration. This retrieval process is rapid, giving rise to savings, and is long-lasting, as it is retained even after one year. However, what is retrieved appears to be simply an aiming direction that enhances behavior on top of an inescapable underlying SPE-based learning process; older individuals in particular do not acquire a general-purpose aiming strategy they can use to complete other adaptation tasks. Thus patients with spinocerebellar ataxia exhibit poor performance in adaptation tasks because they, like healthy individuals, are unable to spontaneously substitute a TE-based aiming strategy for an SPE-based learning system (even when that SPE-based system is impaired), despite having an experimentally-demonstrable latent capacity to do so.

## ACKNOWLEDGEMENTS

We would like to thank the Johns Hopkins Ataxia Center for their assistance and support in patient recruitment. In addition, we thank Alexander D. Forrence for his assistance in data collection for the adaptation task. Finally, we also thank Mitchell Slapik, Jordan Mandel, and Ryan Bloes for cognitive data collection and consensus ratings.

## Abbreviations

SPE: sensory-prediction error
TE: target error

## REFERENCES

Anderson BA, Laurent PA, Yantis S. Value-driven attentional capture. Proc Natl Acad Sci U S A 2011; 108(25): 10367–71.

Anguera JA, Bernard JA, Jaeggi SM, Buschkuehl M, Benson BL, Jennett S, et al. The effects of working memory resource depletion and training on sensorimotor adaptation. Behav Brain Res 2012; 228(1): 107–15.

Anguera JA, Reuter-Lorenz PA, Willingham DT, Seidler RD. Contributions of spatial working memory to visuomotor learning. J Cogn Neurosci 2010; 22(9): 1917–30.

Bastian AJ. Learning to predict the future: the cerebellum adapts feedforward movement control. Curr Opin Neurobiol 2006; 16(6): 645–9.

Bates D, Maechler M, Bolker B, Walker S. Fitting linear mixed-effects models using lme4. J Stat Softw 2015; 67: 1–48.

Bock O, Girgenrath M. Relationship between sensorimotor adaptation and cognitive functions in younger and older subjects. Exp Brain Res 2006; 169(3): 400–6.

Bond KM, Taylor JA. Flexible explicit but rigid implicit learning in a visuomotor adaptation task. J Neurophysiol 2015; 113(10): 3836–49.

Brembs B. Mushroom bodies regulate habit formation in Drosophila. Curr Biol 2009; 19(16): 1351–5.

Brudner SN, Kethidi N, Graeupner D, Ivry RB, Taylor JA. Delayed feedback during sensorimotor learning selectively disrupts adaptation but not strategy use. J Neurophysiol 2016; 115(3): 1499–511.

Buch ER, Young S, Contreras-Vidal JL. Visuomotor adaptation in normal aging. Learn Mem 2003; 10(1): 55–63.

Butcher PA, Ivry RB, Kuo S-H, Rydz D, Krakauer JW, Taylor JA. The cerebellum does more than sensory-prediction-error-based learning in sensorimotor adaptation tasks. J Neurophysiol 2017; Accepted for publication.

Cashaback JGA, McGregor HR, Mohatarem A, Gribble PL. Dissociating error-based and reinforcement-based loss functions during sensorimotor learning. PLoS Comput Biol 2017; 13(7): e1005623.

Christou AI, Miall RC, McNab F, Galea JM. Individual differences in explicit and implicit visuomotor learning and working memory capacity. Sci Rep 2016; 6: 36633.

Cooper FE, Grube M, Elsegood KJ, Welch JL, Kelly TP, Chinnery PF, et al. The contribution of the cerebellum to cognition in Spinocerebellar Ataxia Type 6. Behav Neurol 2010; 23(1-2): 3–15.

Criscimagna-Hemminger SE, Bastian AJ, Shadmehr R. Size of error affects cerebellar contributions to motor learning. J Neurophysiol 2010; 103(4): 2275–84.

Haith AM, Huberdeau DM, Krakauer JW. The influence of movement preparation time on the expression of visuomotor learning and savings. J Neurosci 2015; 35(13): 5109–17.

Haith AM, Krakauer JW. The multiple effects of practice: skill, habit and reduced cognitive load. Curr Opin Behav Sci 2018; 20: 196–201.

Hegele M, Heuer H. Implicit and explicit components of dual adaptation to visuomotor rotations. Conscious Cogn 2010; 19(4): 906–17.

Hegele M, Heuer H. Age-related variations of visuomotor adaptation result from both the acquisition and the application of explicit knowledge. Psychol Aging 2013; 28(2): 333–9.

Heuer H, Hegele M. Adaptation to visuomotor rotations in younger and older adults. Psychol Aging 2008; 23(1): 190–202.

Hogan MJ, Staff RT, Bunting BP, Murray AD, Ahearn TS, Deary IJ, et al. Cerebellar brain volume accounts for variance in cognitive performance in older adults. Cortex 2011; 47(4): 441– 50.

Hothorn T, Bretz F, Westfall P. Simultaneous inference in general parametric models. Biom J 2008; 50(3): 346–63.

Huang J, Gegenfurtner KR, Schutz AC, Billino J. Age effects on saccadic adaptation: Evidence from different paradigms reveals specific vulnerabilities. J Vis 2017; 17(6): 9.

Huang VS, Haith A, Mazzoni P, Krakauer JW. Rethinking motor learning and savings in adaptation paradigms: model-free memory for successful actions combines with internal models. Neuron 2011; 70(4): 787–801.

Huberdeau DM, Haith AM, Krakauer JW. Formation of a long-term memory for visuomotor adaptation following only a few trials of practice. J Neurophysiol 2015a; 114(2): 969–77.

Huberdeau DM, Krakauer JW, Haith AM. Dual-process decomposition in human sensorimotor adaptation. Curr Opin Neurobiol 2015b; 33: 71–7.

Huberdeau DM, Krakauer JW, Haith AM. Practice induces a qualitative change in the memory representation for visuomotor learning. BioRxiv 2017.

Joiner WM, Smith MA. Long-term retention explained by a model of short-term learning in the adaptive control of reaching. J Neurophysiol 2008; 100(5): 2948–55.

Kim HE, Morehead JR, Parvin DE, Moazzezi R, Ivry RB. Invariant errors reveal limitations in motor correction rather than constraints on error sensitivity. Commun Biol 2018; 1(1): 19.

Krakauer JW, Ghez C, Ghilardi MF. Adaptation to visuomotor transformations: consolidation, interference, and forgetting. J Neurosci 2005; 25(2): 473–8.

Lemaire P. Cognitive strategy variations during aging. Curr Dir Psychol Sci 2010; 19(6): 363–9.

Leow LA, Gunn R, Marinovic W, Carroll TJ. Estimating the implicit component of visuomotor rotation learning by constraining movement preparation time. J Neurophysiol 2017: jn 00834 2016.

Mandolesi L, Leggio MG, Graziano A, Neri P, Petrosini L. Cerebellar contribution to spatial event processing: involvement in procedural and working memory components. Eur J Neurosci 2001; 14(12): 2011–22.

Martin TA, Keating JG, Goodkin HP, Bastian AJ, Thach WT. Throwing while looking through prisms. I. Focal olivocerebellar lesions impair adaptation. Brain 1996; 119 (Pt 4): 1183–98.

Maschke M, Gomez CM, Ebner TJ, Konczak J. Hereditary cerebellar ataxia progressively impairs force adaptation during goal-directed arm movements. J Neurophysiol 2004; 91(1): 230– 8.

Mazzoni P, Krakauer JW. An implicit plan overrides an explicit strategy during visuomotor adaptation. J Neurosci 2006; 26(14): 3642–5.

McDougle SD, Taylor JA. Dissociable roles for working memory in sensorimotor learning. bioRxiv 2018: 290189.

McNay EC, Willingham DB. Deficit in learning of a motor skill requiring strategy, but not of perceptuomotor recalibration, with aging. Learn Mem 1998; 4(5): 411–20.

Morehead JR, Qasim SE, Crossley MJ, Ivry R. Savings upon Re-Aiming in Visuomotor Adaptation. J Neurosci 2015; 35(42): 14386–96.

Morehead JR, Taylor JA, Parvin DE, Ivry RB. Characteristics of Implicit Sensorimotor Adaptation Revealed by Task-irrelevant Clamped Feedback. J Cogn Neurosci 2017; 29(6): 1061– 74.

Rabe K, Livne O, Gizewski ER, Aurich V, Beck A, Timmann D, et al. Adaptation to visuomotor rotation and force field perturbation is correlated to different brain areas in patients with cerebellar degeneration. J Neurophysiol 2009; 101(4): 1961–71.

Reuter-Lorenz PA, Sylvester C-YC. The cognitive neuroscience of working memory and aging. In: Cabeza R, Nyberg L, Park D, editors. Cognitive neuroscience of aging: Linking cognitive and cerebral aging Oxford: Oxford University Press; 2005. p. 186–217.

Schmahmann JD, Sherman JC. The cerebellar cognitive affective syndrome. Brain 1998; 121 (Pt 4): 561–79.

Schween R, Hegele M. Feedback delay attenuates implicit but facilitates explicit adjustments to a visuomotor rotation. Neurobiol Learn Mem 2017; 140: 124–33.

Seidler RD. Differential effects of age on sequence learning and sensorimotor adaptation. Brain Res Bull 2006; 70(4-6): 337–46.

Shmuelof L, Huang VS, Haith AM, Delnicki RJ, Mazzoni P, Krakauer JW. Overcoming motor “forgetting” through reinforcement of learned actions. J Neurosci 2012; 32(42): 14617–21.

Smith MA, Ghazizadeh A, Shadmehr R. Interacting adaptive processes with different timescales underlie short-term motor learning. PLoS Biol 2006; 4(6): e179.

Smith MA, Shadmehr R. Intact ability to learn internal models of arm dynamics in Huntington’s disease but not cerebellar degeneration. J Neurophysiol 2005; 93(5): 2809–21.

Suenaga M, Kawai Y, Watanabe H, Atsuta N, Ito M, Tanaka F, et al. Cognitive impairment in spinocerebellar ataxia type 6. J Neurol Neurosurg Psychiatry 2008; 79(5): 496–9.

Taylor JA, Ivry RB. Flexible cognitive strategies during motor learning. PLoS Comput Biol 2011; 7(3): e1001096.

Taylor JA, Klemfuss NM, Ivry RB. An explicit strategy prevails when the cerebellum fails to compute movement errors. Cerebellum 2010; 9(4): 580–6.

Taylor JA, Krakauer JW, Ivry RB. Explicit and implicit contributions to learning in a sensorimotor adaptation task. J Neurosci 2014; 34(8): 3023–32.

Trewartha KM, Garcia A, Wolpert DM, Flanagan JR. Fast but fleeting: adaptive motor learning processes associated with aging and cognitive decline. J Neurosci 2014; 34(40): 13411–21.

Trouillas P, Takayanagi T, Hallett M, Currier RD, Subramony SH, Wessel K, et al. International Cooperative Ataxia Rating Scale for pharmacological assessment of the cerebellar syndrome. The Ataxia Neuropharmacology Committee of the World Federation of Neurology. J Neurol Sci 1997; 145(2): 205–11.

Tseng YW, Diedrichsen J, Krakauer JW, Shadmehr R, Bastian AJ. Sensory prediction errors drive cerebellum-dependent adaptation of reaching. J Neurophysiol 2007; 98(1): 54–62.

Uresti-Cabrera LA, Diaz R, Vaca-Palomares I, Fernandez-Ruiz J. The effect of spatial working memory deterioration on strategic visuomotor learning across aging. Behav Neurol 2015; 2015: 512617.

Vandevoorde K, Orban de Xivry JJ. Motor adaptation but not internal model recalibration declines with aging. BioRxiv 2018: 292250.

Vaswani PA, Shmuelof L, Haith AM, Delnicki RJ, Huang VS, Mazzoni P, et al. Persistent residual errors in motor adaptation tasks: reversion to baseline and exploratory escape. J Neurosci 2015; 35(17): 6969–77.

Weiner MJ, Hallett M, Funkenstein HH. Adaptation to lateral displacement of vision in patients with lesions of the central nervous system. Neurology 1983; 33(6): 766–72.

Werner S, Bock O. Effects of variable practice and declarative knowledge on sensorimotor adaptation to rotated visual feedback. Exp Brain Res 2007; 178(4): 554–9.

Wolpe N, Ingram JN, Tsvetano KA, Henson RN, Kievit RA, Wolpert DM, et al. Motor learning decline with age is related to differences in the explicit memory system. BioRxiv 2018: 353870.

Wong AL, Shelhamer M. Sensorimotor adaptation error signals are derived from realistic predictions of movement outcomes. J Neurophysiol 2011; 105(3): 1130–40.

